# Single-Walled Carbon Nanotube Probes for Protease Characterization Directly in Cell-Free Expression Reactions

**DOI:** 10.1101/2025.01.11.632549

**Authors:** Sepehr Hejazi, Ryan Godin, Vito Jurasic, Nigel F Reuel

## Abstract

Proteins can be rapidly prototyped with cell-free expression (CFE) but in most cases there is a lack of probes or assays to measure their function directly in the cell lysate, thereby limiting the throughput of these screens. Increased throughput is needed to build standardized, sequence to function data sets to feed machine learning guided protein optimization. Herein, we describe the use of fluorescent single-walled carbon nanotubes (SWCNT) as effective probes for measuring protease activity directly in cell-free lysate. Substrate proteins were conjugated to carboxymethyl cellulose-wrapped SWCNT, yielding stable and sensitive probes for protease detection with a detection limit of 6.4 ng/mL for bacterial protease from *Streptomyces griseus*. These probes successfully measured subtilisin activity in unpurified CFE reactions, surpassing commercial assays. Furthermore, they enabled continuous monitoring of activity during synthesis of subtilisin in both purified and lysate-based CFE systems without compromising protein expression. Surface passivation techniques, such as pre-incubation with cell lysate and supplement components, reduced the initial signal loss and improved probe signal stability in the complex cell lysate environment. These modular probes can be used, as described, for high-throughput screening and optimization of proteases and, with the change of conjugated substrate, a wider range of other hydrolases.

## Introduction

Machine learning-guided directed evolution (MLDE) platforms provide powerful alternatives to traditional natural protein evolution methods,^1,2^ overcoming limitations such as restricted sequence space, which can limit the exploration of multi-site linked mutations across the protein’s sequence.^3^ MLDE platforms also enable the incorporation of both positive and negative feedback from mutations, providing a more comprehensive understanding of sequence-function relationships compared to selection-based approaches that focus solely on top-performing sequences.^4^ Low-N models have recently emerged as a solution to overcome the large dataset requirements of traditional protein engineering approaches.^5^ These models enable efficient exploration of protein sequence space with smaller experimental data sets by leveraging techniques such as active learning ^3^ and Bayesian ^6^ optimization to enhance predictive accuracy with limited input. In-vitro protein synthesis systems (cell-free expression, CFE)^7–10^ can provide the throughput necessary to prototype proteins through the iterative design process,^11–13^ however, there is still a limitation with screening function, especially directly in cell lysate; fluorescent nanoprobes built from single-walled carbon nanotubes (SWCNT) can address this need (Figure 1a).

**Figure 1.**
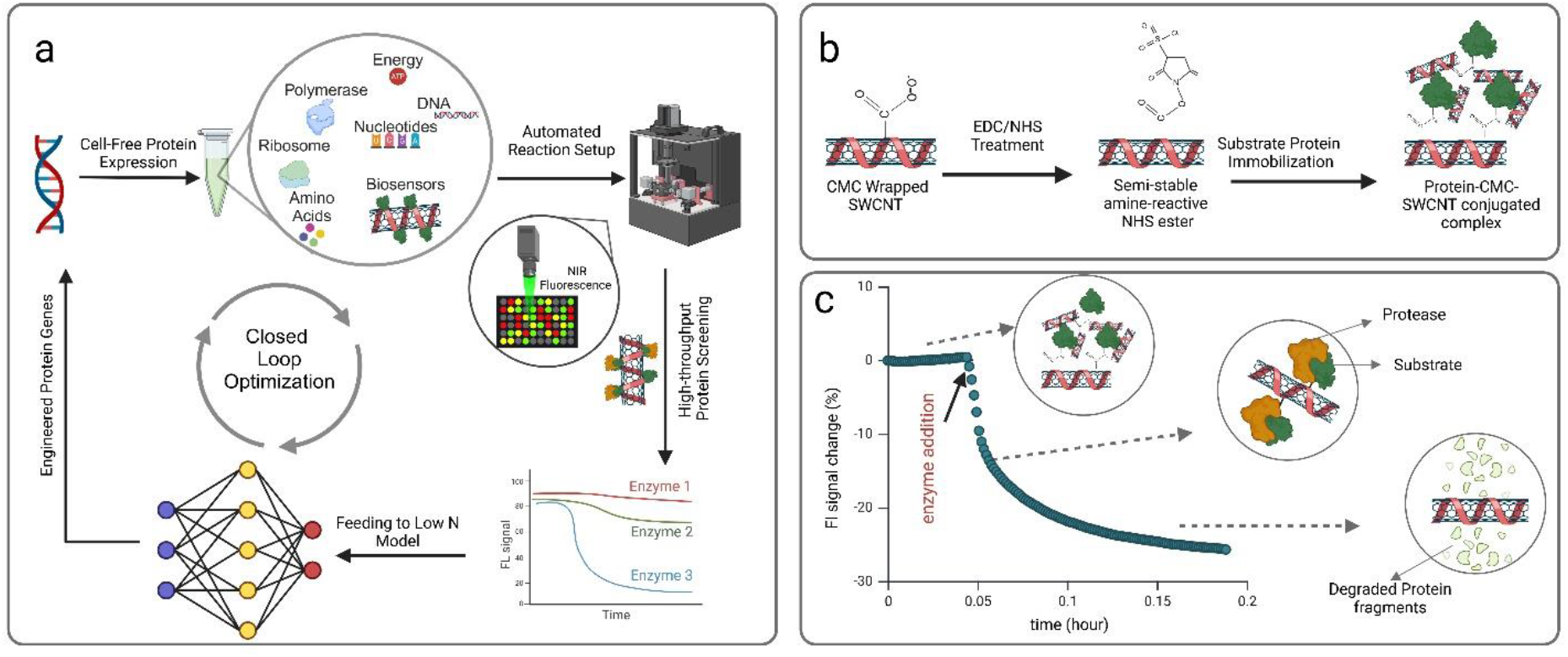
General workflow for Cell-free Expression (CFE)-based protein optimization and utility of single-walled carbon nanotube (SWCNT)-based optical probes. (a) Closed-loop protein optimization with CFE: DNA templates are transcribed and translated using CFE, the proteins are monitored continuously or with endpoint measurements using SWCNT-based optical probes, and their functions are benchmarked. Low-N model learning algorithms are then applied to generate new gene sequences, initiating subsequent cycles of protein design to enhance the target protein. b) Bio-conjugation strategy for substrate immobilization onto CMC-wrapped SWCNT. The process begins with the activation of carboxyl groups on CMC using EDC/NHS chemistry. This is followed by the conjugation of the protein substrate through its surface amine groups (lysine residues), resulting in the formation of a protein-CMC-SWCNT complex. c) Enzymatic reaction between protein-SWCNT probes and the target protease. The initial fluorescence intensity baseline of the SWCNT probes is established before introducing the target enzyme into the solution. Changes in the fluorescence signal are monitored over time, capturing the rate of protein substrate degradation by the protease.

Traditional methods for screening protein function directly in lysate are hampered by the complex lysate expression environment. For proteases, conventional methods like byproduct characterization using liquid chromatography are difficult to resolve against background molecules and are slower, labor-intensive, and limited to periodic or endpoint measurements, highlighting the need for real-time (continuous) monitoring.^14^ Substrate immobilized assays (heterogeneous) such as surface-enhanced Raman spectroscopy,^15^ surface plasmon resonance,^16^ and enzyme-linked protease assays,^17^ offer stability but often suffer from low sensitivity due to solid-liquid enzymatic reaction limitation, high background noise, and complexity in large-scale applications.^18^ Traditional dispersed assays (homogeneous) assays like fluorescence resonance energy transfer (FRET), are simpler to operate and better suited for high-throughput screening but struggle with stability in complex environments, such as cell lysates, and require specialized substrates for each protease.^14,18^ Therefore, novel homogeneous assays based on nanomaterials such as gold nanoclusters^19^, gold nanoparticles^20,21^, and carbon nanotubes^22^ have been developed.

Near-infrared (NIR) fluorescent SWCNT have been used to develop optical sensing probes for the detection of a wide range of biomolecules such as nucleotides,^23^ metabolites,^24^ neurotransmitters,^25^ and enzymes.^26,27^ SWCNT absorb photons from an excitation source, forming excitons, which diffuse along the nanotube and emit fluorescent light upon recombination.^28^ Several properties of SWCNT make them particularly suitable for sensing cell-free expression (CFE) products directly in the reaction mixture. These include their exceptional photostability,^29^ fluorescence in the NIR-II window, which minimizes autofluorescence and scattering in biological samples,^30^ and *in vivo* biocompatibility.^31,32^ SWCNT-based sensors have been used to measure protein binding using cell-free expressed proteins that had a his-tag to immobilize them near the nanotube surface.^33^ This prior CFE and SWCNT work was used as an endpoint assay and was not capable of monitoring protein expression continuously in the lysate reaction, nor was it well suited for measuring enzyme activity.

Established methods for enzyme detection using single-walled carbon nanotubes (SWCNT) involve the non-covalent functionalization of nanotubes with target substrates through direct tip-sonication^34^ or wrapping exchange.^35^ This produces substrate-wrapped SWCNT probes that respond to the degradation of the wrapped substrate through increased solvent access and signal quenching. They have been used to screen biomass and biofilm extracellular polymeric substance (EPS) degrading hydrolases^27,36^. However, this approach relies on the interaction between the hydrophobic regions of the substrate and the SWCNT, which restricts its applicability to molecules capable of forming stable SWCNT dispersions. ^37,38^ In the case of protein substrates, only a limited number, such as bovine serum albumin (BSA) and lysozyme, can achieve this stability. To address these limitations, our group developed a hybrid method that combines covalent bonding of the protein substrate with a nanotube’s non-covalent wrapping, utilizing carbodiimide conjugation chemist to form amide bonds between the free amines of substrate proteins and the carboxyl groups of carboxymethyl cellulose (CMC)-wrapped SWCNT.^39^

In this study, we first investigated the impact of SWCNT probes on protein expression within the cell-free expression CFE system. Next, we used protein-conjugated CMC-SWCNT probes to quantify the endpoint proteolytic activity of CFE products from various proteases and different expression levels of specific proteases. Additionally, continuous monitoring of protease synthesis and activation during CFE was successfully achieved in both lysate-based and purified systems using conjugated probes and conventional protein-wrapped SWCNT probes. Finally, we explored passivation strategies to enhance the stability and sensitivity of these sensors in CFE systems for continuous monitoring.

## Materials and Methods

### SWCNT probes preparation

All chemicals were obtained from Sigma-Aldrich unless otherwise specified.

#### Protein-wrapped SWCNT probes

2mg/ml of Lysozyme or BSA proteins were dissolved in 10mM Tris-HCl buffers with different adjusted pHs (pH 7 to 10) and 10 mM ammonium acetate buffers (pH 4 to 6). Then 1 mg/ml of CoMoCAT (6,5)-enriched single-walled carbon nanotubes (SWCNT; CHASM SG65i, diameter = 0.78 nm) were added. The mixture was sonicated using a Qsonica CL-18 tip sonicator at 6–7 W power (typically 3-4 ml volume of solution) for 90 minutes while placed in an ice bath. The resulting CMC-SWCNT suspension was centrifuged at 14,000 × g for 30 minutes, and 80% of the supernatant was carefully collected to avoid low-stability SWCNT.

#### Conjugated probes

We modified our previous protocol that was used for ECM proteins immobilization.^39^ Carboxymethyl cellulose (CMC; MW = 700 kDa, degree of substitution = 0.8) was dissolved in deionized (DI) water, achieving a concentration of 2 mg/mL. Higher CMC concentrations were avoided to prevent the gelation of the solution. SWCNT were added to achieve a final concentration of 1 mg/mL. The mixture was sonicated at 6–7 W power (typically 3-4 ml volume of solution) for 60 minutes while placed in an ice bath. The resulting CMC-SWCNT suspension was centrifuged at 14,000 × g for 30 minutes, and 80% of the supernatant was carefully collected to avoid low-dispersion SWCNT.

#### Carboxyl group activation of CMC-SWCNT

To adjust the pH to approximately 5, 10% (V/V) μl of 0.1 M MES buffer (pH = 5) was added to the SWCNT suspension. 1-ethyl-3-(3-dimethylaminopropyl)carbodiimide (EDC) (Thermo Scientific) and Sulfo-N-hydroxy succinimide (Sulfo-NHS) (Thermo Scientific) were dissolved in DI water to concentration of 50 mg/ml, then added to the SWCNT solution to reach final concentrations of 4 mg/ml for EDC and 10 mg/ml for Sulfo-NHS, maintaining the same molar ratio. The mixture was vortexed briefly and incubated at room temperature for 1 hour on an orbital shaker with 180 RPM. To quench the EDC activation reaction, 20 mM final concentration of 2-mercaptoethanol was added, and also MES buffer was neutralized by adding 1 M NaOH according to the MES buffer concentration used.

#### Protein Immobilization

To prepare the solution, SWCNT were dispersed in 1X PBS buffer (pH 7.4) to achieve a final concentration of 0.5 mg/ml. Separate solutions of BSA (50 mg/ml) and casein (20 mg/ml) were prepared by dissolving BSA powder in water and casein in an alkaline NaOH solution (1 mM, pH 10), respectively. The proteins were then added into the SWCNT dispersion at a final concentration of 2 mg/ml and incubated overnight at 4°C.

#### Washing and Resuspension

A hydroxylamine solution (10 mM in 1X PBS) was employed as a washing and storage buffer to terminate the immobilization process by providing excess amine groups. After 16 hours of incubation, the protein-SWCNT complex aggregates were collected via centrifugation at 12,000 RCF for 10 minutes at the tube’s bottom. To expedite aggregation, 10–15 μl of 1M HCl was optionally added. The supernatant was discarded, and 1 ml of the washing buffer was used to resuspend the complexes through pipetting. This washing step was repeated three times, after which the samples were stored in 1–2 ml of washing buffer at 4°C. For enhanced baseline stability, mild sonication (5 minutes at 0.5 W) was applied to resuspend the probes, though pipetting alone sufficed for simpler resuspension.

#### SWCNT Concentration Measurements

The concentration of SWCNT were determined spectrophotometrically by measuring absorbance at 661 nm. For this, 100 μl of the SWCNT solution was analyzed in a 96-well plate and compared to a 1 mg/ml SWCNT stock solution as a reference standard.

### Enzymatic reaction and fluorescence measurement

SWCNT NIR fluorescence intensity signals were measured using a custom-built NIR fluorometer (Supporting figure S1), as described in our previous works ^26,27^. The SWCNT probes were diluted 100-to 300-fold before the reaction, ensuring steady fluorescence signals between 6–9 mV as determined by the fluorometer. The diluted probes were then distributed into 96-well plates for purified enzyme experiments and 384-well plates for cell-free expression (CFE) assays.

Protease from *Streptomyces griseus* and Proteinase K were prepared following the manufacturers’ recommended preparation and activation protocols. Varying concentrations of the enzymes were added to the SWCNT probes, and fluorescence signals were recorded. The relative signal change was calculated using the following equation:

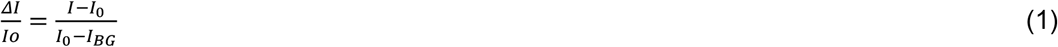

Where *I* is the instantaneous signal, *I*_*0*_ is the signal at time 0, and *I*_*BG*_ is the signal from an empty well.

For spectral scanning measurements, a MiniTracer spectrophotometer (Applied NanoFluorescence, Texas, USA) equipped with a 638 nm laser excitation source was used. Emission spectra were recorded in the range of 900–1400 nm.

### Cell lysate preparation

Cell lysate for cell-free expression reactions were prepared from the One Shot™ BL21 Star™ (DE3) strain of *E. coli* (Thermo Fisher Scientific). Cells were grown and harvested according to a previously published protocol, ^40^ then flash frozen in liquid nitrogen and stored at -80°C until further processing was performed. To prepare lysate from harvested cells, the cell pellet was thawed on ice, re-suspended by vortexing in 1 mL of cold S30 acetate buffer (10 mM tris-acetate pH 8.2, 14 mM magnesium acetate, and 60 mM potassium acetate) per gram of cell mass, and lysed in an Avestin EmulsiFlex-B15 homogenizer at 24,000-26,000 psi. Following a previous report,^41^ the lysed cells were collected and centrifuged at 12,000 xg and 4°C for 10 min. The supernatant was then collected and re-centrifuged at 12,000 xg and 4°C for 10 min. At this point, the resulting lysate supernatant was collected, flash frozen in aliquots in liquid nitrogen, and stored at -80°C for use in cell-free expression reactions.

### Cell-free expression reactions

For lysate-based assays, we used an energy mix (master mix) formulation described in our previous research^40,42^. The cell-free reaction was initiated by combining the following components: cell lysate (24% v/v), energy mix (33% v/v), 5 nM DNA template, and water to adjust the final reaction volume to 15 μL per well. For reactions involving SWCNT probes, 2.5 μl of the water component (50% of the required water volume) was replaced with the SWCNT solution. The reaction mixtures were incubated either on a heated platform of a fluorometer or in an incubator at 30°C to start the reaction.

For PURExpress^®^ (New England BioLabs, Ipswich, MA) assays, the reaction mixture consisted of 40% Solution A, 30% Solution B, 2.25 μL of SWCNT solution, 150 ng of DNA template, and water to adjust the final volume to 15 μL. These reactions were incubated on a fluorometer’s heated platform at 37°C.

### Passivating SWCNT probes with polymers

The passivation strategy reported in prior research by Gilkwad et al. was adapted for the passivation of our SWCNT probes.^43^ Briefly, Poly-L-Lysine (PLK) and polyethylene imine (PEI) polymers were incubated with the SWCNT probes at a 25:1 mass ratio of polymer to SWCNT for 2 hours at 4°C. For the pre-incubation strategy with CFE components, 2.5 μL of non-passivated sensors were added to the CFE reaction and pre-incubated at 4°C for 2 hours prior to the addition of DNA templates. Subsequently, the CFE reaction was initiated by adding the DNA templates and incubating at 30°C. To assess the success of passivation, 2.5 μL of polymer-passivated sensors were added to the CFE reaction, and the initial fluorescence drop during the first hour was measured. Additionally, the expressed reporter protein, sfGFP, fluorescence signal was quantified after 8 hours of the CFE reaction.

## Results and Discussion

### Probe preparation and sensor design

Direct sonication of SWCNT with protease target proteins such as BSA and lysozyme was first attempted as a benchmark for sensor performance in CFE. This direct sonication approach has been used to disperse nanotubes for varied applications over the past few decades^44^, including our first work on hydrolase measurement. Because the protein-SWCNT complex relies on hydrophobic interactions^45^, the probes are inherently unstable, and only a few proteins are able to successfully disperse SWCNT in water with suitable shelf-lives (>1 week); also the resulting dispersions are highly sensitive to environmental factors like pH and ionic strength, limiting the sensor’s functionality in complex environments such as cell-free expression (CFE) systems.

Successful suspensions were achieved for BSA-SWCNT complexes at pH values ranging from 4 to 10, using 10 mM ammonium acetate for acidic conditions and 10mM Tris buffer for basic conditions, and for lysozyme-SWCNT complexes from pH 4 to 9 in the same buffers similar to previous findings^27,46^. Notably, neither protein could suspend SWCNT in 1x PBS at pH 7.4, likely due to the higher ionic strength of the buffer. Another downside of the direct wrapping approach is likely damage to the protein structure due to the suspension protocol that typically needs more than 30 minutes of sonication.^47^ Directly sonicated probes exhibited poor performance in both endpoint and continuous monitoring of lysate-based CFE protease activity (Supporting Information figures S2–S3). Lysozyme-wrapped probes were unable to differentiate the signal from subtilisin-expressing samples and sfGFP controls. While BSA-wrapped probes successfully distinguished endpoint responses for different genes (Supporting Information, Figure S2b), they exhibited instability in lysate-based CFE reactions and failed to differentiate subtilisin expression from sfGFP or no-DNA controls (Supporting Information, Figure S3).

To address the limitations of the direct-wrapping approach, an alternative dispersion method was used to conjugate the target substrate protein to a polymer-wrapped SWCNT. The free surface amine groups of the protein substrates (from exposed Lysines) are conjugated to the amine-reactive NHS ester groups on carboxy methyl cellulose (CMC) wrapped SWCNT that have carboxyl groups activated by EDC^48^ (**Figure 1b**). The presence of multiple free primary amine sites on the protein substrates and the abundance of carboxylic groups on the CMC wrapping facilitate the formation of a cross-linked protein-CMC-SWCNT complex. The formation of this complex also serves as the basis for washing and purifying protein-SWCNT probes, as optimized in our previous work.^39^ This process enhances the sensitivity of hydrolysis biosensors by removing unbound substrate proteins and residual chemicals from earlier steps. While the washing step is primarily employed for insoluble proteins due to aggregation after immobilization, in the case of BSA, lowering the pH with hydrochloric acid can induce aggregation, enabling further purification of the probes.

To confirm the successful conjugation of BSA and casein to CMC-SWCNT, FTIR scans of CMC-SWCNT before and after protein immobilization were performed (Supporting Information, Figure S1). Strong peaks were observed around 1647 cm^−1^ for casein-CMC-SWCNT and 1653 cm^−1^ for BSA-CMC-SWCNT, indicating the presence of amide bonds suggesting the incorporation of casein and BSA onto the probes. ^49,50^ These purified NHS-EDC mediated conjugate sensors with improved stability and reduced background are what we then used to study applicability to prototyping in CFE.

### Sensor characterization and enzymatic response

We next analyzed the response of protein-conjugated probes to a commercial bacterial protease cocktail, specifically bacterial protease from *Streptomyces griseus*. The emission spectrum of BSA- and casein-functionalized probes were recorded before and 30 minutes after exposure to the bacterial protease (Supporting Information, Figure S4). A reduction in fluorescence intensity was observed across the emission spectrum, indicating enzymatic activity, though minimal peak shifts were detected, with the primary changes attributed to intensity variations. This response is linked to substrate degradation, which increases SWCNT access to the polar solvent, resulting in a reduction in the fluorescence intensity.^36^

To quantify the sensor’s response to varying protease concentrations, fluorescence intensity changes over time were monitored using a fluorometer, as shown in **Figure 2b** for casein-CMC-SWCNT probes reacting with *Streptomyces griseus* protease. Among the probes tested, casein sensors exhibited greater response and also better sensitivity based on their LOD of 6.4 ng/ml compared to the BSA-based sensor LOD of 320 ng/ml (Supporting Information, Figure S5). A logarithmic fit of the 30-minute fluorescence intensity response to enzyme concentration showed a strong correlation, with an R-squared value of 0.993 (**Figure 2c**), which will be further used to calculate the equivalent concentrations for different proteases expressed in CFE.

**Figure 2.**
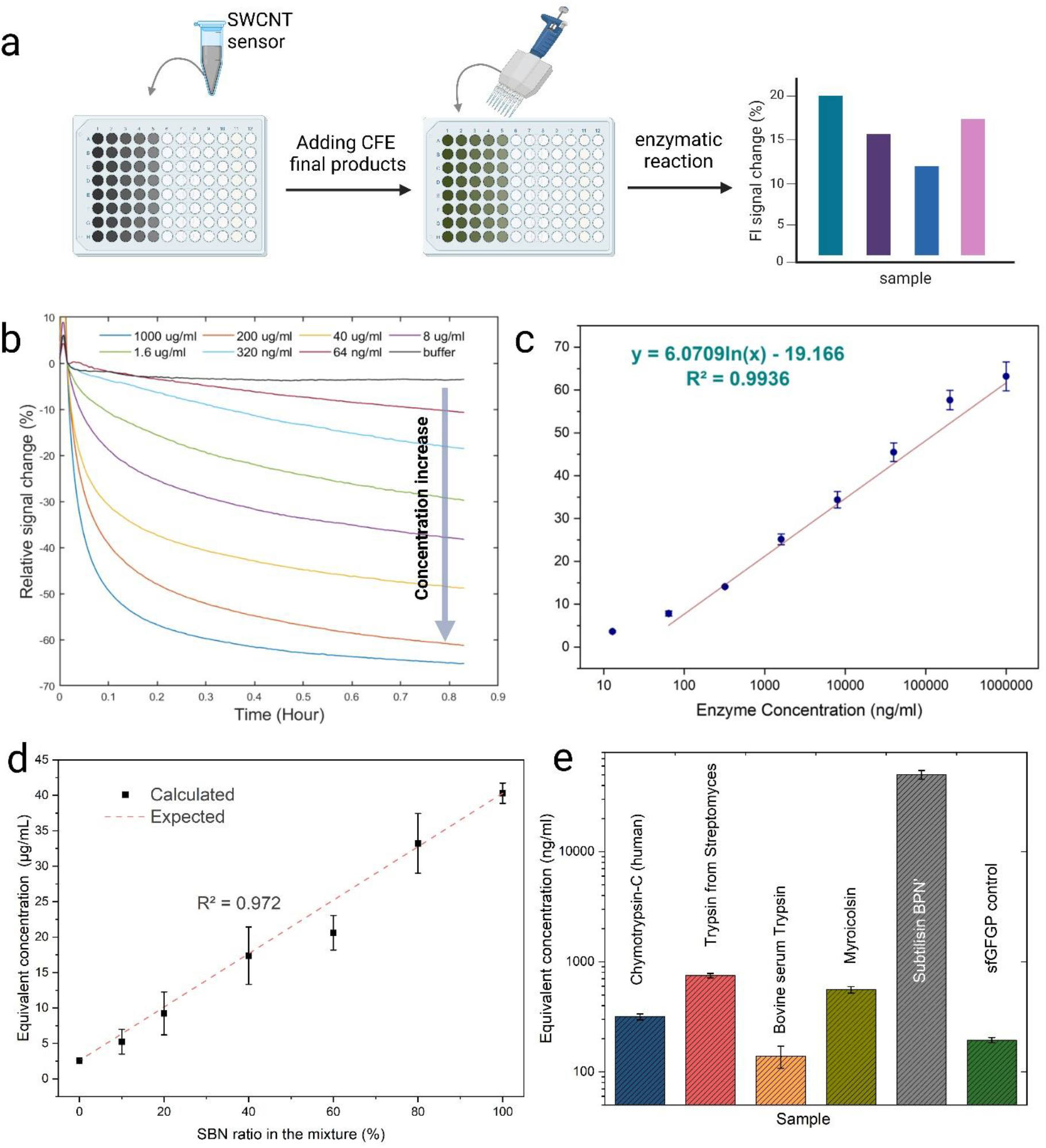
Endpoint enzyme quantification using protein-SWCNT probes (a) Schematic representation of endpoint enzyme concentration measurement using protein-SWCNT probes. The unpurified CFE products are added to the probes, and fluorescence changes in the SWCNT are measured as the output signal. (b) Response of casein-CMC-SWCNT probes to varying concentrations of bacterial protease from *Streptomyces griseus*. (c) Standard curve for the 30-minute response of bacterial protease on casein-CMC-SWCNT probes, with logarithmic fitting and corresponding R-squared value. (d) Endpoint calculated concentrations of enzymes (expressed as bacterial protease equivalents based on sensor signal change) for different mixtures of subtilisin BPN’ CFE and sfGFP control at known dilution ratios exhibiting expected linear parity. (e) Calculated bacterial protease equivalent concentrations for unpurified CFE products expressing protease genes, measured using casein-CMC-SWCNT probes.

### Testing CFE endpoint enzyme activity without purification

To evaluate the capability of conjugated and directly wrapped probes for detecting protease hydrolysis activity without purification, subtilisin BPN’, an extracellular alkaline serine protease known for robust expression in CFE systems,^51^ was expressed alongside sfGFP as a control to verify successful protein expression. The resulting products were added to the probes, and changes in SWCNT fluorescence intensity over time were monitored. Casein-conjugated probes demonstrated the highest response, with a 40% Δ*I*/*I*_0_ decrease after 30 minutes for subtilisin, and exhibited the lowest standard deviation among replicates compared to BSA-conjugated, BSA-wrapped, and lysozyme-wrapped probes (Supporting Figures S7), therefore, casein-CMC-SWCNT probes were selected for quantification in subsequent experiments.

Next, we mixed subtilisin BPN’ and no-DNA CFE endpoint products in various ratios to evaluate the ability of casein-conjugated probes to quantify subtilisin at different expression levels. It is important to note that diluting subtilisin with buffer would not account for the background protease activity of CFE systems; therefore, a mixture with no-DNA controls or inert gene products was used for more accurate measurement. The 30-minute response of the probes to these mixtures was used to calculate the equivalent bacterial protease concentration, which was then plotted against the mixing ratios. A linear relationship was observed, with an R-squared value of 0.972, demonstrating the probes’ ability to distinguish different expression levels of subtilisin in CFE systems without purification (**Figure 2d**).

### Quantification of different proteases expressed in CFE

As a feasibility demonstration of these probes’ capability to differentiate between protease activities, various serine proteases from different species were expressed using a CFE system, and the enzymatic activity of the end products was measured using casein-conjugated probes. The equivalent bacterial protease concentrations from *Streptomyces griseus* were calculated based on the sensor signal response and are presented in **Figure 2e**. Among the tested proteases, the prokaryotic genes for trypsin from *Streptomyces griseus* and Myroicolsin from *Myroides profundi D25* exhibited the highest activity, while eukaryotic genes of chymotrypsin-C from humans and bovine serum trypsin showed the lowest activity. It is important to note that none of the four genes expressed were optimized for our CFE system prior to this study, and the optimal hydrolysis reaction condition for each enzyme is different but they were tested in the same condition. Also, at this stage of development, these ranked activities are on a per lysate volume normalization not a per protein level. The measured activity is influenced by the specific activity of the protease as well as the expression level of the gene, which can vary widely depending on the sequence used. The expressed proteins could be purified, quantified, and then assayed to get specific activities (normalized by amount of protein), but this would obviate the HTS nature of the current approach. A few methods for in-lysate quantification have been recently presented, such as split GFP and tetracysteine tag use,^52–55^ but were not applied in this current study. Instead, the focus is on the functionality of the probes for in-lysate measurements of the enzymes; forthcoming work from our group will show the approach and importance of normalization, especially when generating standardized training sets for machine learning optimization approaches.

### Effect of SWCNT probes on the CFE protein expression

To assess the impact of SWCNT probes on protein expression in a CFE system, Casein and BSA conjugated probe solutions were used to replace the nuclease-free water portion of the CFE reaction. We replaced 100% (5 µl) or 50% (2.5 µl) of the water with SWCNT solution (1mg/ml SWCNT concentration) and measured sfGFP fluorescence intensity after 10 hours of expression as an indicator for successful expression (**Figure 3b**). The fluorescence intensities were normalized to the no-SWCNT control (mean = 1, SD = 0.0658). The 50% addition of BSA-conjugated (mean = 0.987, SD = 0.0398) and casein-conjugated (mean = 0.939, SD = 0.0321) SWCNT solutions had minimal impact on sfGFP expression. However, increasing the volume of SWCNT solution to 100% (replacing water) decreased normalized fluorescence intensity for both BSA-conjugated (mean = 0.829, SD = 0.0479) and casein-conjugated (mean = 0.851, SD = 0.0327) probes. Therefore, 50% SWCNT solution replacement to the free water volume of the CFE reaction was used in all subsequent experiments.

**Figure 3.**
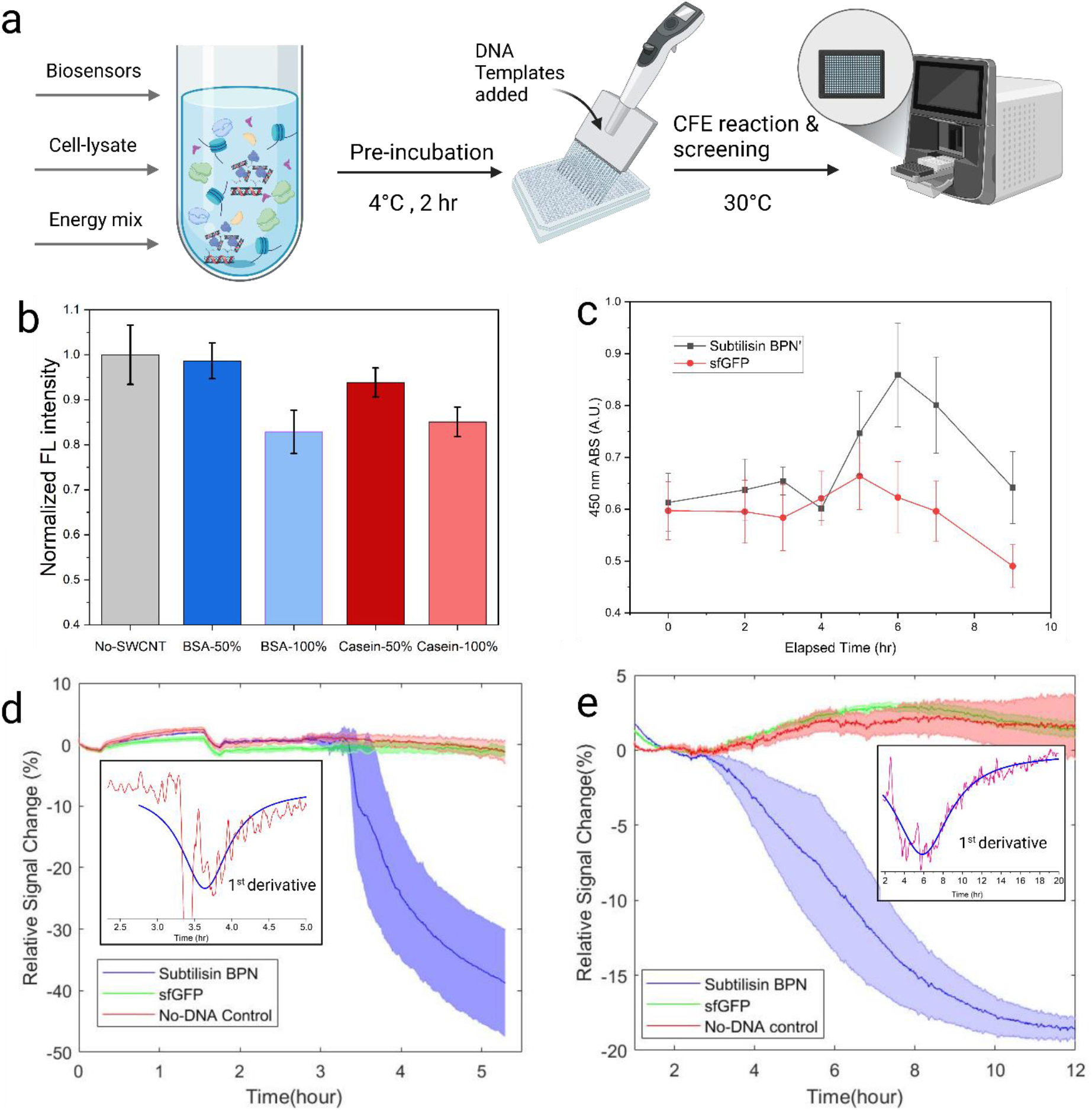
Continuous measurement of enzyme expression in the CFE system using protein-SWCNT probes. (a) Schematic of the dynamic measurement setup and experimental process. (b) Effect of adding BSA and c asein-SWCNT probes to the CFE system on sfGFP protein expression, with different volumes of SWCNT probe solution replacing water in the reaction. (c) Absorbance measurements from the Pierce colorimetric protease activity assay at different time points for SBN and sfGFP expression. Averages of three replicates for each gene are shown as bars with standard deviation as error bars. (d) Response of directly wrapped BSA-SWCNT probes in the PureExpress CFE system with genes templates added at time = 0 h and (e) Response of BSA-CMC-SWCNT probes in-lysate-based CFE systems with gene templates added at time = 0 h, both showing the average of three replicates as solid lines with standard deviation as the shaded region.

### Dynamic measurements of protease activity inside the CFE systems

One of the key advantages of using SWCNT probes is their ability to provide continuous responses to environmental changes, offering valuable information such as expression time and activity over time in CFE systems. We leveraged this capability to use these sensors for dynamic measurements of protease activity in-lysate-based and PureExpress systems. First, we utilized a commercial protease activity assay, the Pierce colorimetric assay, which uses a casein-based substrate to quantify protease activity. Subtilisin and sfGFP genes were expressed in the lysate-based CFE system, and the protease activity of the mixture (without purification) was measured using the Pierce colorimetric assay. Every 1 or 2 hours from the start of the reaction, 2 µL of the reaction mixture was sampled, and the absorbance of the TNBSA dye was measured as a signal for enzymatic activity (**Figure 3c**). Subtilisin activity started to differentiate from the sfGFP control after 5 hours of sampling and continued to increase until the 6-hour measurement, followed by a decrease observed for both sfGFP and subtilisin samples, likely due to hydrolysis of the expressed proteins from the expressed and native proteases present in the lysate.

Next, we used SWCNT-based probes to monitor dynamic protein expression and subtilisin activation in both pure and lysate-based CFE systems. In the PureExpress system, BSA-wrapped SWCNT were employed with subtilisin and sfGFP control genes. The relative signal changes over time are shown in **Figure 3d**, revealing a sharp onset of hydrolysis activity approximately 3.3 hours after the reaction began. While BSA-wrapped SWCNT successfully detected enzymatic activity in the PureExpress system, they proved ineffective in the lysate-based CFE system, likely due to the latter’s higher ionic strength and the presence of more complex biomolecules (Supporting Information figure S3).

To address this, we utilized casein- and BSA-conjugated probes for dynamic measurements in the lysate-based system. Although casein-CMC-SWCNT performed well for endpoint measurements, they exhibited greater initial instability (30-40% reduction in the 1^st^ hour) during dynamic measurements, likely due to interference from native lysate proteases (Supporting Information figure S9). By contrast, BSA-CMC-SWCNT provided a more stable response in the lysate-based system. **Figure 3e** shows the BSA-CMC-SWCNT response along with the first derivative of the subtilisin signal. The fluorescence change represents an accumulative signal, whereas the slope of the reduction indicates relative enzyme activity at any time point. From the derivative data, we see an increase in activity at the 4-hour mark and maximum activity at 6 hours after the reaction began, as indicated by the derivative minima. This timing aligns with the maximum activity observed using the Pierce colorimetric assay but obtained with much less technician time and experimental materials.

### Surface passivation to reduce the effect of cell-free reaction components on sensor signal response

Although the conjugated probes successfully demonstrated enzymatic activity in a lysate-based assay, they exhibited a significant initial signal reduction (10-20%), particularly during the first hour of the cell-free expression (CFE) reaction (Supporting Information, Figure S10). This reduction was attributed to the complex reaction environment in-lysate-based CFE, specifically the high salt concentration in the energy mix and the presence of native proteins from the lysate. To address this issue, the probes were pre-incubated with the CFE components for 2 hours at 4°C before the addition of the DNA template and subsequent incubation at 30°C to initiate the CFE reaction.

In addition to this approach, the stability of the conjugated probes was further evaluated using two commonly used passivation polymers, Poly-L-Lysine (PLK) and polyethyleneimine (PEI). These polymers have been previously reported to successfully passivate the surface of SWCNT probes.^43^The passivation process involved pre-incubating the probes with the polymers for 4 hours at 4°C, using a 25:1 mass ratio of passivation reagent to SWCNT.

The impact of passivated probes on the cell-free expression level was assessed by adding different probes to a CFE system containing the sfGFP gene and measuring the expressed reporter protein signal after 10 h of reaction (**Figure 4b**). A negative effect on sfGFP production was observed for both BSA- and casein-conjugated probes that underwent passivation with polymers. Probes passivated with PLK and PEI significantly reduced the expressed sfGFP signal, indicating their negative impact on the CFE system. This reduction is likely attributed to interactions of the high concentrations of these positively charged polymers with the transcription or translation machinery components of the CFE system. However, pre-incubation with CFE components did not result in a significant reduction in sfGFP production compared to untreated probes.

**Figure 4.**
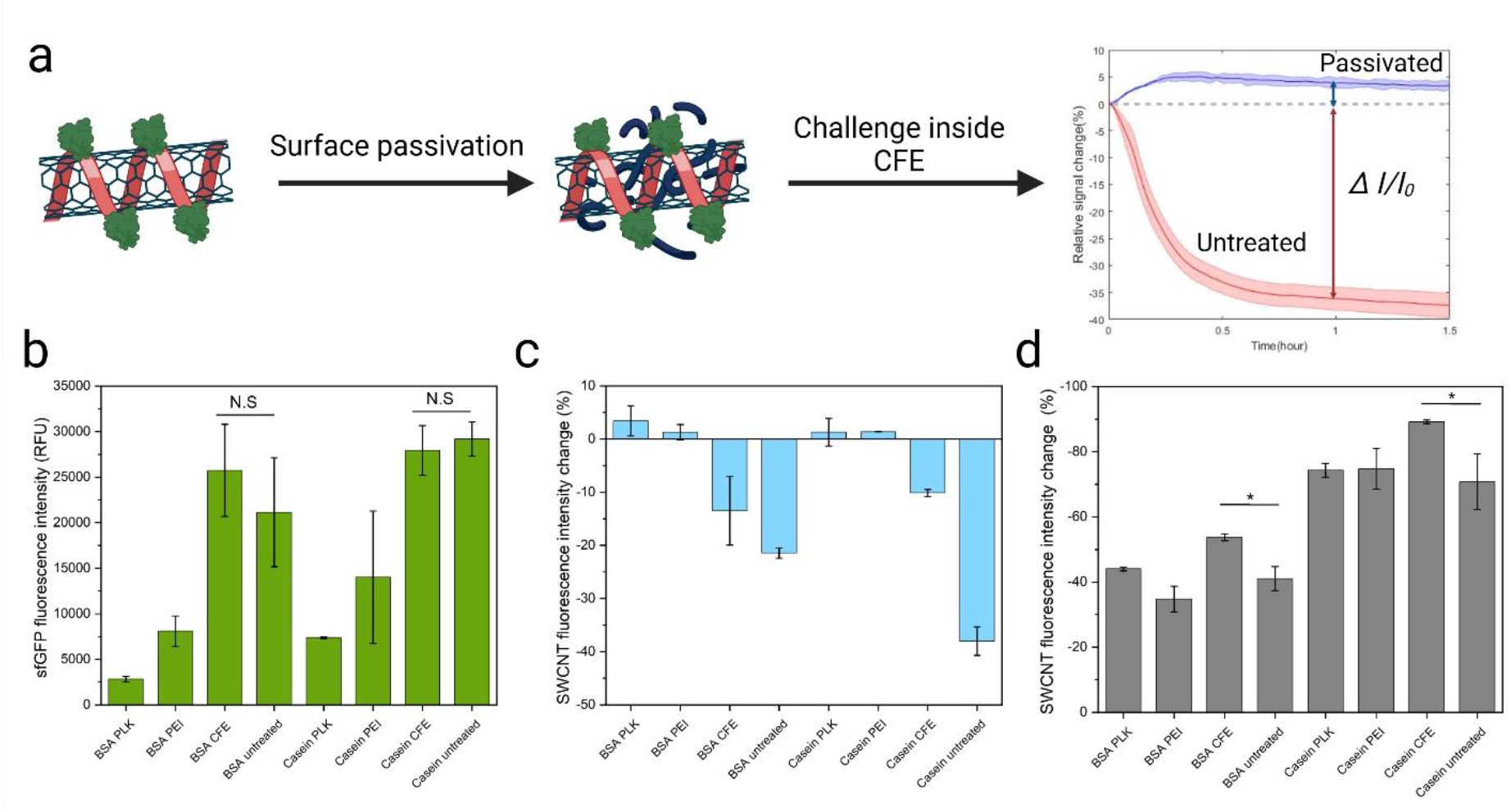
Protein-SWCNT probe surface passivation strategies. (a) Schematic of the passivation process and subsequent testing of the probes within a CFE system. (b) Effect of passivated protein-SWCNT probes on sfGFP fluorescence expression levels in CFE systems. (c) Background signal changes of protein-SWCNT probes after 1-hour incubation in a lysate-based CFE system. (d) Sensor response of passivated and unpassivated protein-SWCNT probes after a 30-minute reaction with 50 µg/mL of proteinase K from *Streptomyces griseus*.

Next, we measured the initial signal drop of the probes during the first hour of the CFE reaction under different passivation strategies. The polymer passivation strategy not only mitigated the initial signal drop but also slightly increased the brightness of the probes after 1 hour in the CFE reaction. Additionally, the CFE component pre-incubation strategy effectively reduced the initial signal drop for BSA-conjugated probes from 21.4% to 13.5% and for casein-conjugated probes from 38.0% to 10.15% (**Figure 4c**).

Finally, the impact of passivation on the sensor response to protease degradation was evaluated by measuring the fluorescence signal of protein-SWCNT probes after a 30-minute reaction with 50 µg/mL of proteinase K from Streptomyces griseus (**Figure 4d**). The PEI and PLK passivation strategies showed no significant impact on the response of BSA- and casein-conjugated probes compared to untreated probes. However, pre-incubation with CFE components significantly increased the protease response of the probes (p-values: 0.0159 for BSA and 0.04612 for casein), likely due to lysate native proteins adsorption onto the SWCNT. This adsorption may have acted as an additional substrate for the protease, thereby enhancing the signal.

## Conclusions

In this study, we successfully developed and tested a modular assay using SWCNT-based probes to monitor protease activity in CFE systems. The conjugation approach enabled immobilization of substrate proteins onto carboxymethyl cellulose (CMC)-wrapped SWCNT probes, generating fluorescence signals upon substrate hydrolysis. These probes demonstrated functionality not only in simple buffer systems but also in complex environments like lysate-based CFE reactions.

Casein-CMC-SWCNT probes were used for endpoint quantification of subtilisin activity across varying expression levels, achieved by mixing the CFE products without purification—which was not possible with the commercial assay tested. Additionally, BSA conjugated probes enabled continuous monitoring of protease expression and activation in-lysate-based CFE systems, providing valuable information about target protein expression and activity. It should be noted that these dynamic measurements were performed using a common 384-well plate, allowing for simultaneous screening of fluorescent protein quantification tags such as split GFP or halo tags, thereby enabling normalization on a per-protein basis. This platform highlights the potential of SWCNT fluorescent probes for high-throughput screening (HTS) of protease activity to provide standardized data for ML-guided protein design. Furthermore, the modularity of the conjugation approach enables immobilization of both high- and low-solubility substrates, facilitating the design of HTS probes for other hydrolases.

Passivation strategies to mitigate the initial fluorescence signal drop in-lysate-based CFE were explored. Although polymer-based passivation with polyethyleneimine (PEI) and poly-L-lysine (PLK) effectively reduced initial signal drop, they negatively impacted protein expression yields in CFE. Pre-incubation with CFE reaction components, while less effective at reducing signal drop, maintained protein expression yields and offered a practical compromise. The residual fluorescence reduction can be compensated for by increasing the light source intensity after the drop. A limitation of these probes is the presence of non-specific binding in complex environments; therefore, exploring more sophisticated passivation strategies could improve their performance in such settings.

In summary, this study demonstrates the versatility, sensitivity, and modularity of SWCNT-based fluorescent probes for monitoring protease activity in cell-free expression (CFE) systems, establishing them as powerful tools for both endpoint and continuous protease assays, particularly in high-throughput screening (HTS) for protein engineering. For instance, SWCNT sensors outperformed commercial assays in endpoint benchmarking of subtilisin BPN’, which is critical for closed-loop optimization of proteases to enhance their activity and catalytic efficiency. Furthermore, we demonstrated the importance of continuous monitoring of proteolytic activity during synthesis in CFE. This approach is essential for accurately benchmarking proteases at their peak activity, as we observed a decline in activity over time during synthesis. This data could enable more robust comparative analysis between mutants in HTS campaigns. The continuous data generated can also be applied to stability studies, facilitating the design of more stable proteases such as ones used in industrial processes or as therapies.

