## Supporting information for "Single-Walled Carbon Nanotube Probes for Protease Characterization Directly in Cell-Free Expression Reactions"

**Supplement Section 1: DNA sequences used in this study**

Below are the gene templates used for the CFE expressed proteins. All sequences were optimized for *Escherichia coli* using IDT's codon optimization tool. Each color represents the region according to Addgene pJI1-sfGFP gene for the Plasmid vector and our added sites for making circularized templates:

T7 Promoter – T7 Promoter

RBS – Ribosome binding site

Start – Start codon

Protein Sequence – Gene coding for protein

Stop – Stop codon

T7 Terminator – T7 Terminator

Circularization site – HindIII Digest

Primer Sequences – Primer sequences

Ori – Ori

KanR – KanR

Supplement Sequence 1*:* sfGFP LET (999 bp)

GTAAAACGACGGCCAGTAGCGCTATTAAAGCTTCGAAATTAATACGACTCACTATAGGGAGACCACAA
CGGTTTCCCTCTAGAAATAATTTTGTTTAACTTTAAGAAGGAGATATACATATGAGCAAAGGTGAAGAAC
TGTTTACCGGCGTTGTGCCGATTCTGGTGGAACTGGATGGCGATGTGAACGGTCACAAATTCAGCGT
GCGTGGTGAAGGTGAAGGCGATGCCACGATTGGCAAACTGACGCTGAAATTTATCTGCACCACCGGC
AAACTGCCGGTGCCGTGGCCGACGCTGGTGACCACCCTGACCTATGGCGTTCAGTGTTTTAGTCGC
TATCCGGATCACATGAAACGTCACGATTTCTTTAAATCTGCAATGCCGGAAGGCTATGTGCAGGAACG
TACGATTAGCTTTAAAGATGATGGCAAATATAAAACGCGCGCCGTTGTGAAATTTGAAGGCGATACCCT
GGTGAACCGCATTGAACTGAAAGGCACGGATTTTAAAGAAGATGGCAATATCCTGGGCCATAAACTGG
AATACAACTTTAATAGCCATAATGTTTATATTACGGCGGATAAACAGAAAAATGGCATCAAAGCGAATTTT
ACCGTTCGCCATAACGTTGAAGATGGCAGTGTGCAGCTGGCAGATCATTATCAGCAGAATACCCCGAT
TGGTGATGGTCCGGTGCTGCTGCCGGATAATCATTATCTGAGCACGCAGACCGTTCTGTCTAAAGATC
CGAACGAAAAACGGGACCACATGGTTCTGCACGAATATGTGAATGCGGCAGGTATTACGTGGAGCCA
TCCGCAGTTCGAAAAATAATAAGTCGACCGGCTGCTAACAAAGCCCGAAAGGAAGCTGAGTTGGCTG
CTGCCACCGCTGAGCAATAACTAGCATAACCCCTTGGGGCCTCTAAACGGGTCTTGAGGGGTTTTTT
GCTGAAAGCGAGACTAAGCTTTAAACTTCGGGTCATA GCTGTTTCCTG

Supplement Sequence 2 : Subtilisin BPN' (1344 bp)

GTAAAACGACGGCCAGTAGCGCTATTAAAGCTTCGAAATTAATACGACTCACTATAGGGAGACCACA
ACGGTTTCCCTCTAGAAATAATTTTGTTTAACTTTAAGAAGGAGATATACATATGGCAGGTAAAAGTAA
CGGCGAGAAAAAATACATCGTTGGCTTCAAACAAACGATGTCGACCATGAGCGCAGCGAAAAAGAAA

GATGTCATCAGCGAAAAAGGCGGTAAAGTGCAGAAACAATTCAAATACGTTGACGCGGCCAGTGCC
ACCCTGAATGAAAAAGCAGTGAAAGAACTGAAGAAAGATCCGTCCGTGGCGTACGTTGAAGAAGAC
CATGTTGCTCACGCGTATGCCCAGTCCGTTCCGTACGGTGTCTCACAAATTAAAGCACCGGCTCTGC
ATTCGCAGGGCTATACCGGTAGCAACGTTAAAGTCGCGGTGATTGATAGCGGCATCGACAGTTCCC
ACCCGGATCTGAAAGTTGCGGGCGGTGCCAGCATGGTGCCGAGCGAAACCAATCCGTTCCAGGAC
AACAATAGCCATGGCACGCATGTGGCGGGTACCGTTGCAGCTCTGAACAATTCTATTGGCGTCCTG
GGTGTGGCACCGTCTGCTAGTCTGTATGCGGTTAAAGTCCTGGGCGCCGATGGCTCTGGCCAGTAC
AGTTGGATTATCAACGGTATTGAATGGGCGATCGCCAACAATATGGATGTGATCAATATGAGCCTGG
GCGGTCCGTCCGGTTCAGCCGCACTGAAAGCAGCTGTCGACAAAGCAGTTGCTTCCGGTGTGGTTG
TTGTGGCCGCAGCCGGTAACGAAGGCACGTCAGGCTCATCGAGCACCGTGGGTTATCCGGGCAAAT
ACCCGTCGGTTATTGCGGTCGGTGCCGTGGATTCTAGTAATCAGCGTGCGAGCTTTTCCTCAGTTGG
CCCGGAACTGGACGTTATGGCCCCGGGTGTCTCTATTCAAAGTACGCTGCCGGGTAACAAATATGG
CGCGTACAATGGTACCAGCATGGCATCACCGCATGTGGCTGGTGCTGCGGCCCTGATCCTGAGCAA
ACACCCGAACTGGACGAATACCCAGGTTCGCTCGAGCCTGGAAAACACCACGACCAAACTGGGCGA
TTCTTTCTATTACGGCAAAGGTCTGATCAATGTTCAGGCAGCTGCGCAATAATAAGTCGACCGGCTG
CTAACAAAGCCCGAAAGGAAGCTGAGTTGGCTGCTGCCACCGCTGAGCAATAACTAGCATAACCCC
TTGGGGCCTCTAAACGGGTCTTGAGGGGTTTTTTGCTGAAAGCGAGACTAAGCTTTAAACTTCGGGT
CATAGCTGTTTCCTG

Supplement Sequence 3 : Bovine serum Trypsin LET (EC:3.4.21.4) (1271 bP)

GTAAAACGACGGCCAGTAGCGCTATTAAAGCTTCGAAATTAATACGACTCACTATAGGGAGACCACAA
CGGTTTCCCTCTAGAAATAATTTTGTTTAACTTTAAGAAGGAGATATACATATGAAGACGTTTATTTTCCT
TGCGTTACTGGGGGCCGCCGTAGCGTTCCCAGTTGATGATGATGACAAAATCGTCGGCGGTTACACA
TGCGGCGCGAATACGGTGCCCTACCAGGTGTCCCTGAACTCGGGCTACCATTTCTGCGGAGGCTCG
TTGATTAACAGCCAATGGGTGGTGTCGGCGGCTCATTGTTACAAATCGGGCATCCAAGTCCGTCTTG
GAGAGGACAACATAAACGTCGTCGAAGGGAACGAACAGTTTATCTCCGCCAGCAAGAGCATCGTTCA
TCCTTCATACAACAGCAATACCTTGAACAACGATATCATGTTAATTAAATTGAAGAGCGCGGCTTCTTTA
AATTCCCGCGTCGCCTCCATTTCGCTGCCAACTTCATGTGCTTCGGCGGGCACGCAGTGTTTAATTAG
CGGATGGGGTAATACGAAATCAAGTGGCACCAGCTACCCCGACGTATTGAAATGCTTGAAAGCTCCTA
TTTTGTCTGACAGCTCCTGCAAGAGCGCATATCCAGGGCAGATTACTTCTAATATGTTCTGTGCAGGTT
ATCTGGAGGGAGGGAAGGATTCATGTCAGGGTGATTCAGGGGGGCCTGTCGTATGCAGCGGTAAGC
TGCAAGGAATAGTCAGTTGGGGCTCGGGTTGCGCGCAAAAGAATAAACCAGGGGTCTACACGAAAGT
CTGCAACTACGTTTCGTGGATTAAGCAGACCATCGCTAGCAACTAATAAGTCGACCGGCTGCTAACAA
AGCCCGAAAGGAAGCTGAGTTGGCTGCTGCCACCGCTGAGCAATAACTAGCATAACCCCTTGGGGC
CTCTAAACGGGTCTTGAGGGGTTTTTTGCTGAAAGCGAGACTAAGCTTTAAACTTCGGGTCATAGCTG
TTTCCTG

Supplement Sequence 4 : Myroicolsin from *Myroides profundi*, H2BKX5 · H2BKX5_MYRPR (2322 bp)

GTAAAACGACGGCCAGTAGCGCTATTAAAGCTTCGAAATTAATACGACTCACTATAGGGAGACCACAA
CGGTTTCCCTCTAGAAATAATTTTGTTTAACTTTAAGAAGGAGATATACATATGCGGAAAATAATCGTTAC
TGGGTTTGCTTTTTTATGCTTGGGAGTGTATGACAATTATGGTCAAACTATATCCACTCGCTCGGCCGT
TAGATCTACCTATGACCTTGACAAGGCAACTAGAGAAATCAATGCATACACCACAAAAGCGGCTAAGAA
CAAACAAGAGGCATTCCTGGAAGCGGAGAAAAGAAATGTACCTATCAGCGGGGTGAACGCGCGCGG
CAACTATTTTGAGCTGTCAGGGATAGACAAAAATGGAATATTACTGTACAAATCAACCCTGAATTATGGC
TCTCGTTTAACTGCCCGTGTGAATGCAATTCAAAAAGAAGTAGGTGTAAACCAATACCTTGAGGGGGA
AGGAATGACGGTGGGTATTATCGATGGTTTACCCTTGTTAGACACGCACCAAGAATTTTATACTACTAC
CTCGAACACCACTAGTCGCGTGACTTTGGGCGAATCCGTGCCAACGTTAACGACTTACAACGCTAAA
GGGTACCAGAAGAGCCGTTTTCATGCTACTCACGTTGGGGCGACCATGGTAGGGTTGGGGTATAACA

ATAAAGCGCAAGGGATTGCCCCTAAGGCTAAACTTGTAAGCTATTCGTGGAATAACGATTACCGCAAG
ATGGGACAGATGGCATCGGGGGGTATGCTGGTGTCCAATCATAGCTATGGGTATAACTACTTCGATGA
TTACGGGTATTTGAACGAGCCTAGTTTGATACGCAATTTTGGAGCATATTCAGAGCACTCTCGGGAGTT
TGATAGAGTGGCATACTTATTTGCGTACTATCAGCCAGTTATAGCAGCCGGCAACGACGGCGAGTACC
ACTACAACGTATATGGTGGGGGTCAAAAGGAGGACTGCAACTGTGATTTATTAAACGATTCATCAGTTT
CGAAGAATGCCGTCGTGGTGGCAGCAGTCGAGCAGGTCGCCCAGTATACGGGACCATCTGACGTCG
TCCTGGCATCTTTTTCTTCTCAAGGCCCTACCAACGACTTCAGAATAAAACCAGATATCTCCGCTAAAG
GAGTCGATGTGTTGTCGGCGGCTTATCGCAATCCTAATCCGTTATATGGTGCTGCGGAAACATCTTTGT
ACGCATATAGCGATGGGACGAGCATGGCGGCTCCAGCGGTTTCGGGCGTCTTCACACTGTGGCAGG
AGTGGGCGATACATGCATCCAGCACGAACATGCCATTTAAATCAGCTACATTGCGTGCCCTGATGGCA
CACACGGCCGATGAAGCAGGCAGAGCAGCGGGCCCAGACCACCTTTTCGGGTGGGGGGTAATTAAT
GCAAAGGCGGGGGTCGAAGTCATGTTGGCAGCTAAGGATAAAAGATCCACATACATTTTGGAAAACG
AGCTTAGAGAACAACAGAAATACACACACGAGATTCAAGTCGGGGAGAAAATGTCTAAAATGGTTGTC
ACTCTTGCCTGGACCGATGCTCCTGGCGTGGTGAGCTACCAAAATAGTGACGAGAATTACAAACGTAA
CAATGGCGACCTTGTCAATGACCTTGATGTGGTTGTTAGAAAAGGCAAAAACACATATTATCCCTGGAT
GTTGAACAAAGATTTCAACGACTTACGGGCGATACAAGGCGTTAATGATGTGGACAACATTGAGAAGA
TAGAATTATATGATGTTGAACCAGGAACGTATGTTATTGAGGTGACGCACAAGGGCAAACTTGAGACC
GGCAAGCAGGAGTATAGCTTAATTTCTACTGTTGGTGAGTTTGATGATCTTCAAGAGTCGAAGGTAGA
AACCAAACAAGAAATACGGCTTTGGCCGAATCCTGTAGAAGATCATTTATATGTTAGTTTGGATAAGAC
GTACAACGGAAAGGTAATTGACATGAAAGTCTACGATATTAATGGACGGCTTGTCCTGTCTTCGTCTGA
AACCGTACAGCAAGAAAAGGTTAGCATAAATATGGCATCCTTAAATTCGAATATCTACGTTGTGGAGGT
GAAAGGTGACAACCTGTCTAAAACTGTACGCATCGCGAAGCGGTAATAAGTCGACCGGCTGCTAACA
AAGCCCGAAAGGAAGCTGAGTTGGCTGCTGCCACCGCTGAGCAATAACTAGCATAACCCCTTGGGG
CCTCTAAACGGGTCTTGAGGGGTTTTTTGCTGAAAGCGAGACTAAGCTTTAAACTTCGGGTCATAGCT
GTTTCCTG

Supplement Sequence 5: Trypsin from *Streptomyces sp*. CB00072 (1068 bp)

GTAAAACGACGGCCAGTAGCGCTATTAAAGCTTCGAAATTAATACGACTCACTATAGGGAGACCACAA
CGGTTTCCCTCTAGAAATAATTTTGTTTAACTTTAAGAAGGAGATATACATATGAAGCACTTTCTGCGTG
CATTAAAAAGATGTTCTGTAGCTGTAGCAGTCGCAACTGTGGCTATTGCGGTTGTTGGTTTACAGCCG
GTAACCGCTAGTGCGGCGCCCAATCCAGTGGTAGGCGGTACTAGAGCAGCTCAAGGAGAGTTCCCG
TTTATGGTCAGACTTAGCATGGGGTGCGGAGGAGCATTATACGCCCAAGATATTGTACTTACGGCTGC
GCATTGCGTCTCGGGTAGCGGTAACAACACTTCGATTACAGCTACTGGGGGGGTCGTCGATTTACAAT
CGCCGAATGCGGTTAAGGTCAGATCGACCAAAGTACTTCAGGCTCCGGGATACAACGGGACGGGCA
AGGACTGGGCTCTTATCAAATTGGCCCAGCCTATTAACCAACCCACATTGAAGATTGCGACTACGACG
GCGCATAACCAAGGGACCTTCACGGTGGCGGGCTGGGGAGCTAATCGTGAAGGGGGGTCCCAACA
ACGCCATTTACTGAAAGCTAATGTACCTTTCGTAAGTGATGCTGCTTGTCGTTCGGCCTACGGGAACG
AACTGGTGGCGAACGAGGAAATATGCGCTGGTTATCCAGATACTGGAGGGGTTGACACTTGTCAAGG
AGACAGTGGAGGTCCAATGTTCCGGAAAGACAACGCTGATGAGTGGGTTCAGGTAGGCATAGTAAGT
TGGGGTTACGGGTGCGCACGTCCAGGCTATCCAGGCGTGTACGCAGAGGTGAGCACGTTTGCGAGT
GCTATCGCCTCCGCTGCTCGCACGTTGTAATAAGTCGACCGGCTGCTAACAAAGCCCGAAAGGAAGC
TGAGTTGGCTGCTGCCACCGCTGAGCAATAACTAGCATAACCCCTTGGGGCCTCTAAACGGGTCTTG
AGGGGTTTTTTGCTGAAAGCGAGACTAAGCTTTAAACTTCGGGTCATAGCTGTTTCCTG

Supplement Sequence 6: Chymotrypsin-C (human) EC:3.4.21.2 (1089 bp)

GTAAAACGACGGCCAGTAGCGCTATTAAAGCTTCGAAATTAATACGACTCACTATAGGGAGACCACAA
CGGTTTCCCTCTAGAAATAATTTTGTTTAACTTTAAGAAGGAGATATACATATGTTGGGGATAACAGTTT
TGGCGGCATTATTGGCCTGCGCTTCATCATGTGGGGTACCGTCTTTCCCACCTAATTTAAGTGCCAGA

GTGGTCGGCGGCGAAGACGCTCGGCCGCATTCATGGCCATGGCAAATCAGCCTGCAATATTTGAAGA
ATGATACTTGGAGACATACATGCGGTGGAACATTGATAGCGTCAAACTTCGTTTTAACGGCAGCCCATT
GTATCTCCAACACGCGTACCTACCGTGTAGCGGTGGGTAAGAATAACCTTGAAGTCGAAGACGAAGA
GGGCAGCCTGTTTGTCGGAGTCGATACTATACACGTTCATAAGCGCTGGAATGCTTTGCTGTTGCGTA
ATGATATTGCCTTAATAAAGTTAGCAGAGCATGTCGAATTGTCGGACACTATACAGGTAGCATGCCTTC
CAGAGAAGGATTCCTTACTTCCCAAGGACTATCCCTGCTATGTTACAGGGTGGGGTCGGCTGTGGAC
TAACGGCCCAATCGCAGATAAGTTGCAGCAAGGATTACAACCTGTGGTAGATCACGCGACTTGCTCTC
GTATAGACTGGTGGGGCTTCAGAGTAAAAAAGACCATGGTTTGTGCTGGCGGGGATGGGGTTATTTC
TGCTTGTAATGGAGATTCGGGTGGACCGTTGAATTGTCAACTTGAGAATGGCTCATGGGAGGTTTTTG
GAATAGTTTCCTTTGGTTCCCGCAGAGGTTGCAACACGCGTAAAAAGCCGGTTGTCTACACTCGCGT
TTCCGCGTATATAGATTGGATCAACGAAAAAATGCAACTGTAATAAGTCGACCGGCTGCTAACAAAGCC
CGAAAGGAAGCTGAGTTGGCTGCTGCCACCGCTGAGCAATAACTAGCATAACCCCTTGGGGCCTCTA
AACGGGTCTTGAGGGGTTTTTTGCTGAAAGCGAGACTAAGCTTTAAACTTCGGGTCATAGCTGTTTCC
TG

**Supplement Section 2: Supplement figures used in this study**

**Schematic and details of the custom built NIR fluorometer used in this study to measure SWCNT emission intensity**

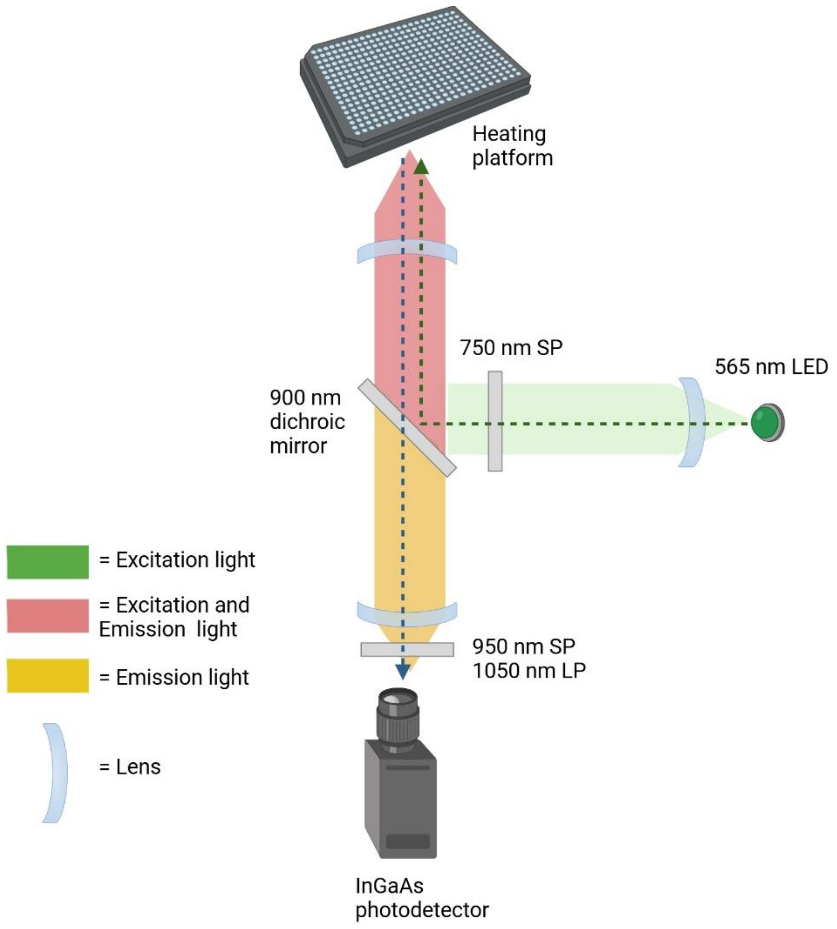

*Figure S1. Schematic of the NIR fluorometer used in this study to measure SWCNT emission intensity.*

**Probe response to CFE proteins with direct sonicated protein wrapped SWCNTs.**

The following figures illustrate the response of protein-wrapped SWCNT probes to end-products of cell-free expression (CFE) (Figure S2). After establishing a baseline, analytes were added to the probe solution at a 5% of the total volume (95% SWCNT solution) . The initial signal drop can be attributed to three factors: dilution of the SWCNT concentration due to the addition of the analyte buffer, intrinsic signal instability of the SWCNTs to the CFE reaction buffers, and the unrecorded fluorescence data during sample addition on the fluorometer.

Figure S3 shows the response of directly sonicated BSA-SWCNT probes within the cell-free reaction. These probes experienced a signal loss of more than 50% and were unable to distinguish between subtilisin and control samples.

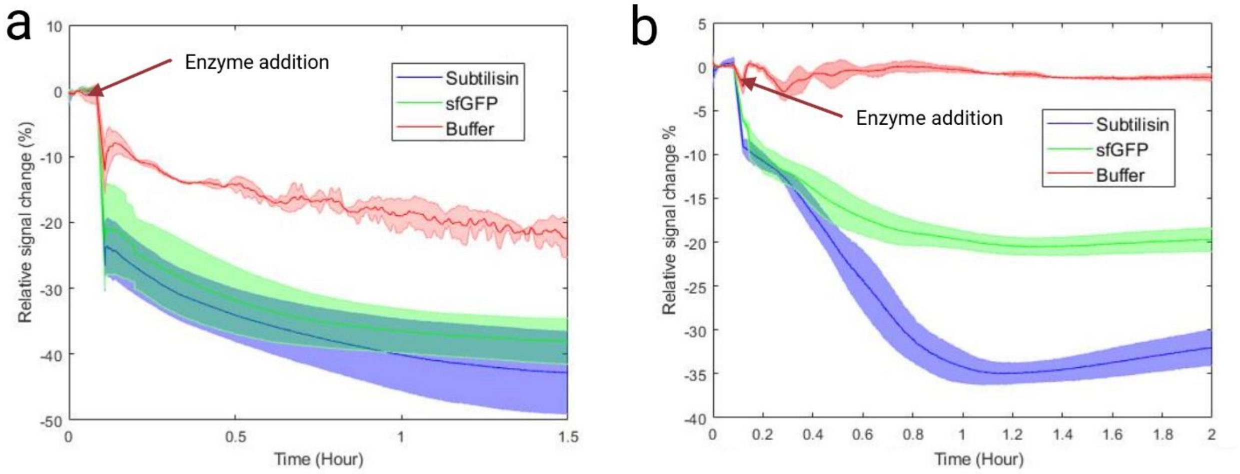

*Figure S2. Probe response to CFE proteins with direct sonicated protein wrapped SWCNTs. a) lysozyme-wrapped SWCNT respond to different CFE products of subtilisin and sfGFP as control. Buffer added to test sensor stability upon small volume addition. b) BSA-wrapped SWCNT sensor response to different CFE products of subtilisin and sfGFP as control*

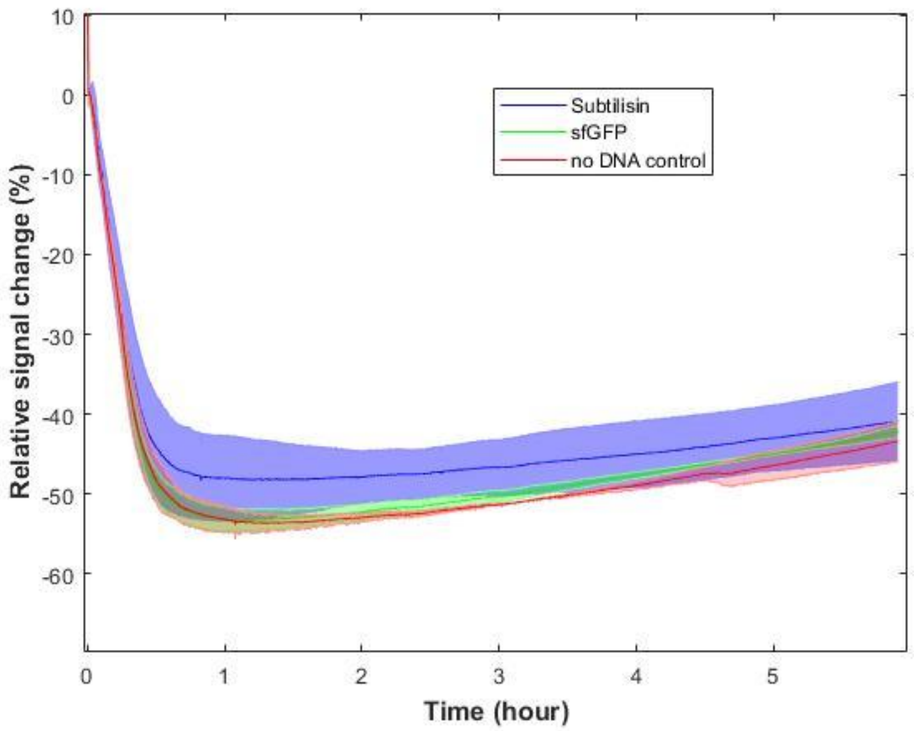

*Figure S3. Direct sonicated BSA-wrapped sensor response inside CFE with different genes. Subtilisin as target, sfGFP to check the expression and no DNA control.*

**Characterization of conjugated protein SWCNT probes**

Figure S4 presents the Fourier Transform Infrared Spectroscopy (FTIR) analysis of CMC-SWCNT, BSA-conjugated, and casein-conjugated probes. Key peaks are labeled, including the amide bond characteristic of protein samples (1647 $cm^{-1}$), highlighting the successful conjugation of proteins to the SWCNTs.

Figure S6 depicts the emission spectral changes of SWCNTs before and after 30 minutes of reaction with bacterial protease. These changes provide insights into the interaction between the SWCNT probes and the protease, demonstrating their potential for detecting enzymatic activity.

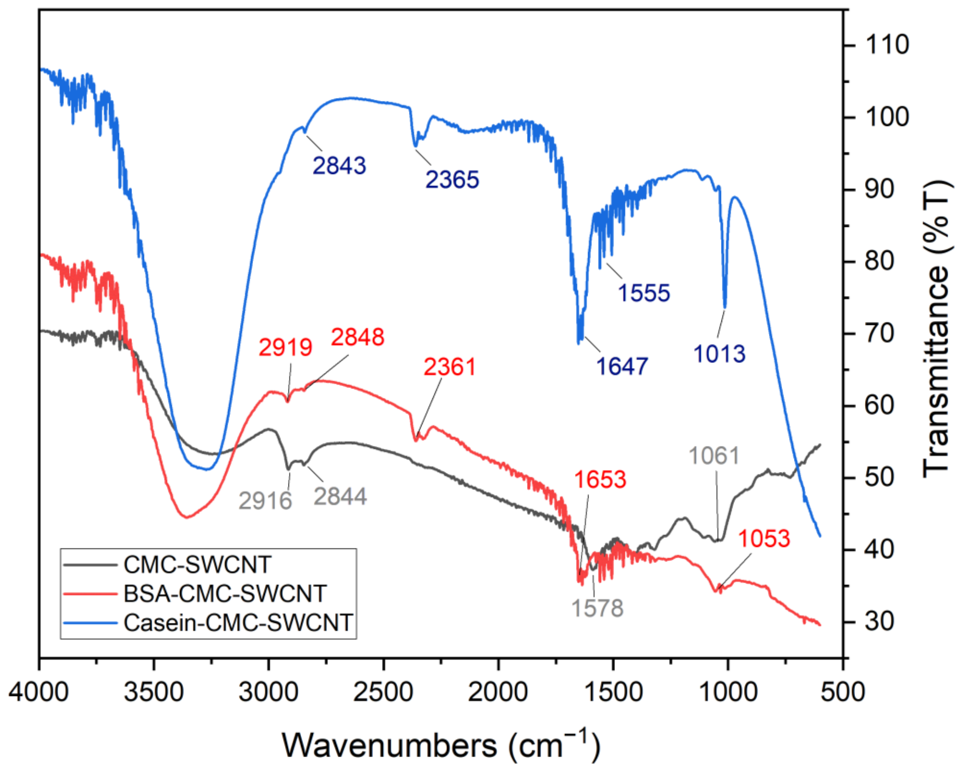

*Figure S4. FTIR plot for the different SWCNT probes with CMC wrapping (black), BSA conjugated CMC-SWCNT (red), and casein conjugated CMC-SWCNT (blue)*

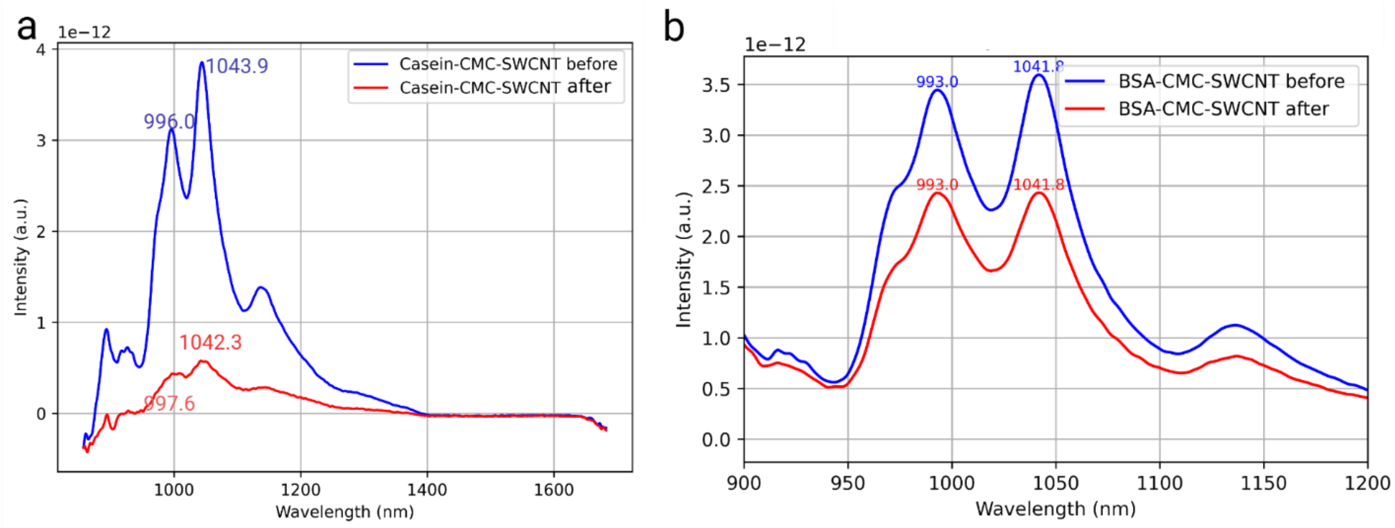

*Figure S5 Emission spectra of a) casein conjugated and b) BSA conjugated probes before and after 30 minutes of incubation with 100 ug/ml of bacterial protease.*

**Conjugated protein SWCNT probes response to proteases**

The following figures (S7,S8) describe the responses of conjugated protein probes to addition of bacterial protease or CFE end products added to the probes. Figure S7 is the BSA conjugated probe response to different levels of bacterial protease. Figure S8 describes the response of Bsa conjugated probes to different expression levels of subtilisin end-products. Figure S9,S10 show the response of casein and BSA conjugated probes inside the lysate based CFE reaction with the genes added from the beginning.

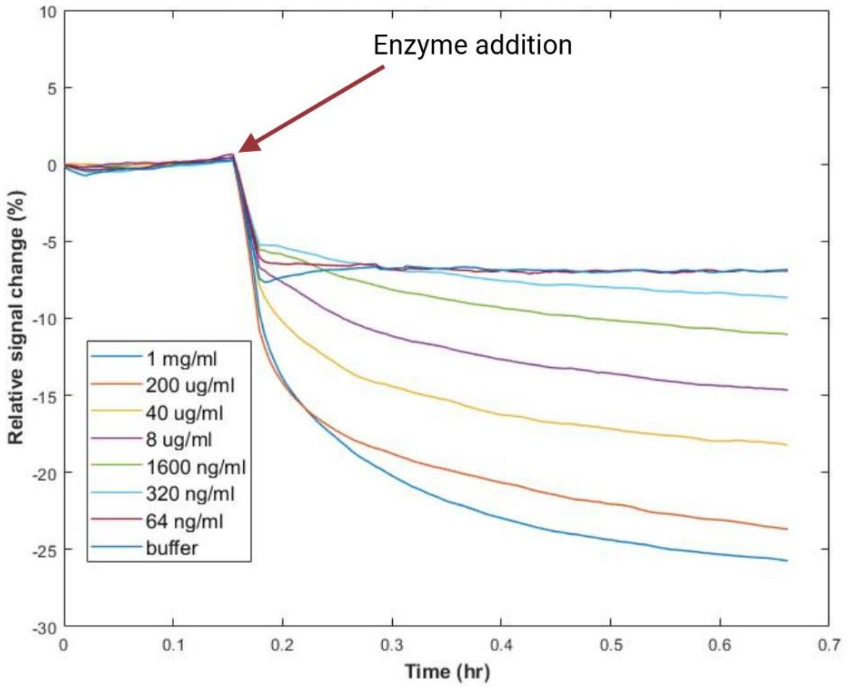

*Figure S7. BSA conjugated sensor response to different concentrations of bacterial protease*

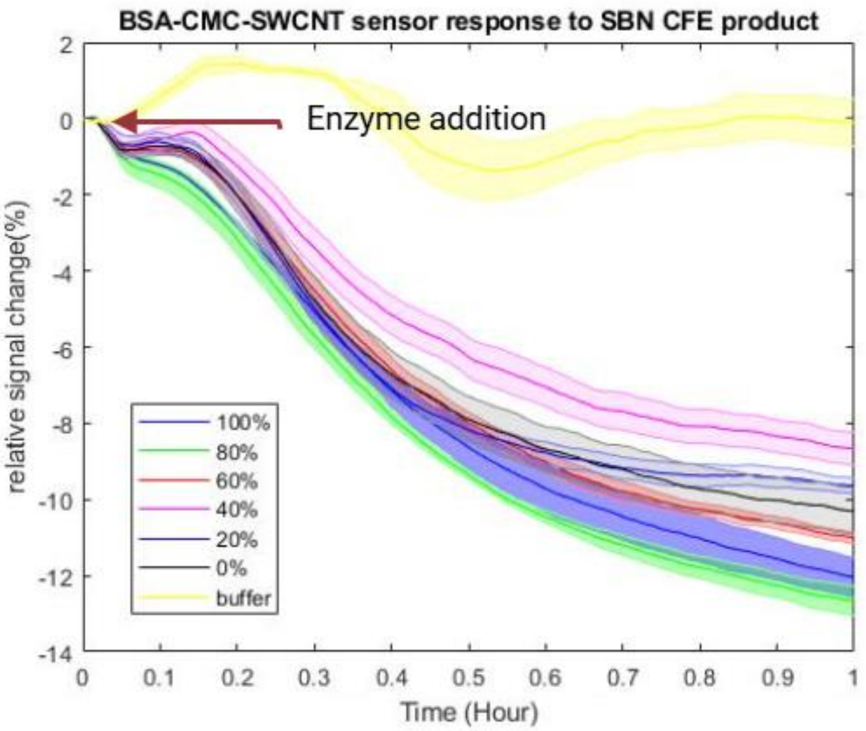

*Figure S8. BSA conjugated sensor response to different mixtures of subtilisin and sfGFP CFE end product mixtures.*

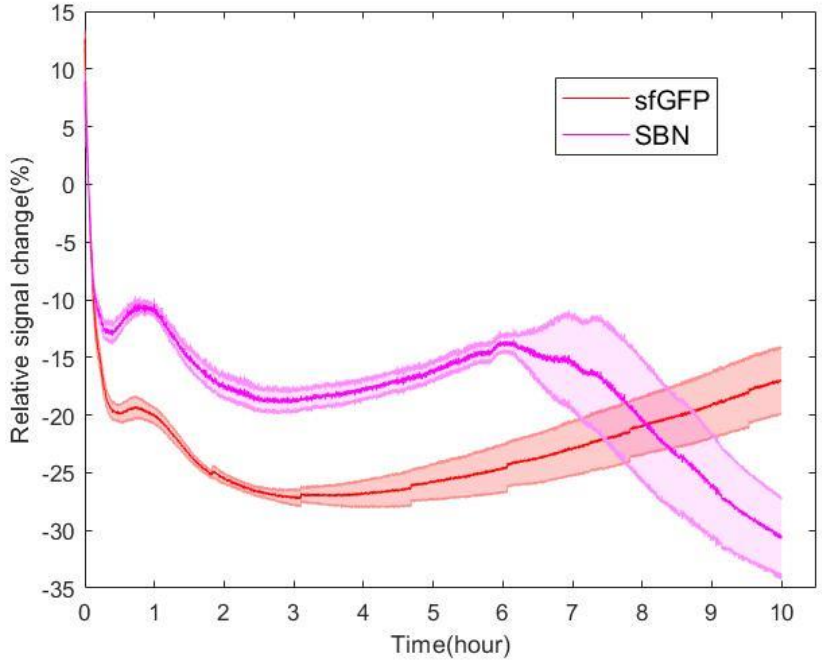

*Figure S9. Casein conjugated probe response in the lysate-based CFE system with genes templates added at time = 0 h, showing the average of three replicates as solid lines with standard deviation as*

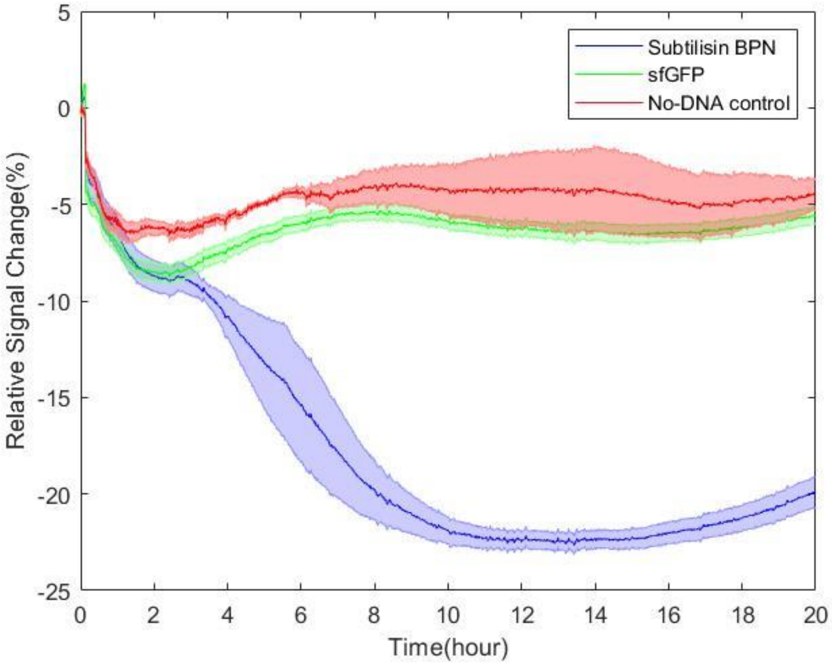

*shaded region.*

*Figure S10. BSA conjugated probe response in the lysate-based CFE system with genes templates added at time = 0 h, showing the average of three replicates as solid lines with standard deviation as shaded region.*
